# Multiple Linear Regression: Bayesian Inference for Distributed and Big Data in the Medical Informatics Platform of the Human Brain Project

**DOI:** 10.1101/242883

**Authors:** Lester Melie-Garcia, Bogdan Draganski, John Ashburner, Ferath Kherif

## Abstract

We propose a Multiple Linear Regression (MLR) methodology for the analysis of distributed and Big Data in the framework of the Medical Informatics Platform (MIP) of the Human Brain Project (HBP). MLR is a very versatile model, and is considered one of the workhorses for estimating dependences between clinical, neuropsychological and neurophysiological variables in the field of neuroimaging. One of the main concepts behind MIP is to federate data, which is stored locally in geographically distributed sites (hospitals, customized databases, etc.) around the world. We restrain from using a unique federation node for two main reasons: first the maintenance of data privacy, and second the efficiency in management of big volumes of data in terms of latency and storage resources needed in the federation node. Considering these conditions and the distributed nature of data, MLR cannot be estimated in the classical way, which raises the necessity of modifications of the standard algorithms. We use the Bayesian formalism that provides the armamentarium necessary to implement the MLR methodology for distributed Big Data. It allows us to account for the heterogeneity of the possible mechanisms that explain data sets across sites expressed through different models of explanatory variables. This approach enables the integration of highly heterogeneous data coming from different subjects and hospitals across the globe. Additionally, it offers general and sophisticated ways, which are extendable to other statistical models, to suit high-dimensional and distributed multimodal data. This work forms part of a series of papers related to the methodological developments embedded in the MIP.

## INTRODUCTION

The amount of neuroimaging data (i.e. MRI, fMRI, PET, EEG, etc.) continues to expand exponentially with the production of thousands of studies worldwide as part of clinical and research activities. As an example, after just a few decades of the Magnetic Resonance Imaging technique (MRI), hundreds of thousands of individuals of all ages and conditions have been scanned and the rate of MRI data collection is set to grow considerably over the coming years. Unfortunately, the majority of this data is locally warehoused in laboratories and hospitals and is therefore poorly capitalized into new knowledge, scientific publications and the development of novel medical therapies. This raises the urgent need for a more efficient exploitation of such limitless richness of information in order to shed light on the principles of brain anatomy and function in both healthy and pathological states. In recent years, in light of this necessity, the neuroscientific and software developer’s communities have joined efforts to embrace new paradigms related to data sharing, virtualization and federation. These paradigms provide a framework to integrate large amounts of data that will undoubtedly help in generating more realistic models of brain function. Data integration is an effective approach devoted to appropriately combining neuropsychological, genetic, clinical and neuroimaging multimodal data (at the micro, meso and macro scales), in a very heterogeneous population coming from different parts of the globe under dissimilar conditions (race, education level, dietary conditions, etc.).

Precisely, the Medical Informatics Platform (MIP) as part of the Human Brain Project (Subproject 8 (SP8)) (https://www.humanbrainproject.eu/en/medicine/medical-informatics-platform/), is conceived to provide an informatics infrastructure to the neuroscientists and clinicians studying brain anatomy and function using clinical along with multimodal big data. This platform faces the challenges of adapting, designing and implementing novel technologies in terms of hardware (clusters etc.), software (i.e. NoSQL, Apache Spark) and data analytics (i.e. decentralized machine learning) to deal with the large and geographically distributed nature of the data. In this paper, we propose a theoretical framework based on the ‘Bayesian formalism’ that contributes to the development of mathematical tools to deal with big and distributed data as part of the MIP environment. Rather than introducing a fully new mathematical development, we gather together and adapt general advances previously described in the Bayesian modeling literature to ultimately propose our methodological framework. We extended these developments to the versatile Multiple Linear Regression (MLR), which is considered one of the workhorses of statistics and data modeling in the neuroimaging field. The Bayesian formalism is ideal for the core of our theoretical machinery since it deals, in a natural way, with the problem of having big, heterogeneous and distributed data.

The paper in general is organized as follows. The main section, ‘Theory’, is divided into two main subsections. The first is devoted to providing the grounds of the general Bayesian framework for distributed data in two ‘regimes’: Parallel and Streaming. A general schema is illustrated that defines the local (hospitals) and federation (global) nodes; the latter federates and delivers the information to the end user. Based on previous works, a model selection/averaging step is proposed, constituting the third level of Bayesian inference. The second subsection describes a special case of the general principles outlined in the former subsection, which are applied to the multiple linear regression model. In addition, a summary of the general equations to be used in practice is shown.

## THEORY

### 1. Problem Statement

One of the main goals of data analysis is to identify the models that best explain the data, along with their intrinsic parameter distributions, in order to discover basic organizational and functional principles of natural systems. Conventional modeling setups and algorithms presuppose that data is stored in one single location as a unique data source (i.e. database). Therefore, new techniques are required to model data when it is large and geographically distributed.

### 2. General Bayesian computational framework: Parallel approach

Figure 1 shows the general scheme of our computational framework for distributed data. Data is dispersed across different remote sites, which in our case are hospitals/databases that may be located in different countries. The upper part of the figure represents the Federation node (aka master node, centralized node) that federates the aggregated information (or aggregates) coming from local nodes or hospitals (aka decentralized nodes, worker nodes) represented in the lower part of the figure. Within the framework of this paper, these aggregates are expectations of various model parameters. In this case, the federated estimation of parameter distributions is obtained using local aggregates provided in parallel by all local nodes. This setting is denominated as ‘parallel’ (Broderick et al., 2013).

**Figure 1.**
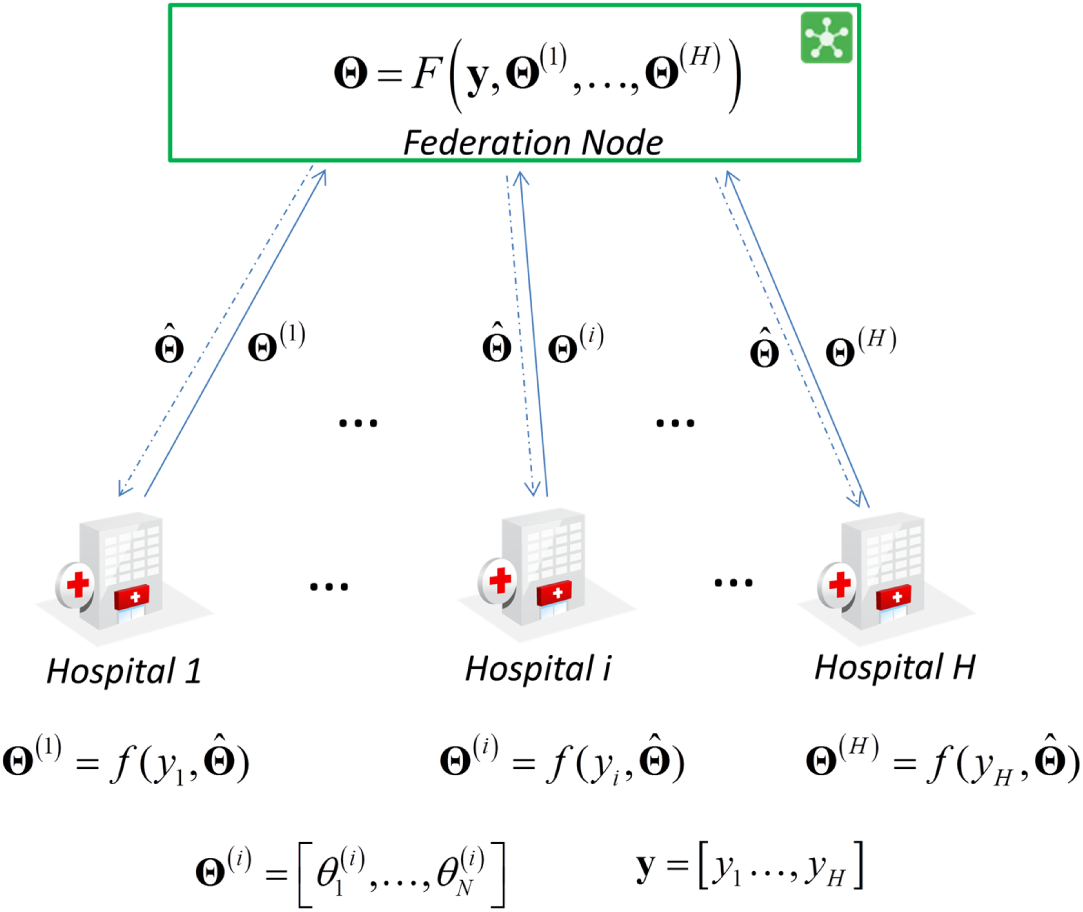
General scheme of the parallel paradigm in the Medical Informatics Platform (Human Brain Project).

In this paradigm, the information can be interchanged in both directions. Within an iterative algorithm, the federation node provides intermediate parameter estimates 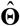 downstream and receives aggregates (**Θ**^(^*^i^*^)^) from each local node. The set of aggregates from Hospital ‘*i*’ is represented by 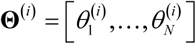 and *y_i_* is the data for the hospital ‘*i*’, where *H* is the number of hospitals so that 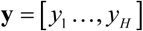.

We proposed the use of the Bayesian formalism as a theoretical framework to define our computational engine, which we named ‘*Bayesian computational framework*’ (BCF). Therefore our BCF is based on the fundamental Bayes theorem so that the estimation of the parameters 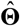 at the federation node is expressed by the following equation:

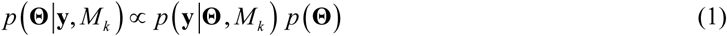

where *p*(**Θ**|**y**,*M_k_*) is the posterior distribution of parameters **Θ** for the specific model *M_k_* and 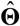 the set of parameter values that maximize 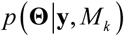; 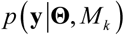 is the likelihood (taking into account data from all Hospitals) and *p*(**Θ**) the prior probability of the parameters of interest.

Assuming that chunks of data belonging to each hospital are independent, the likelihood can be expressed as:

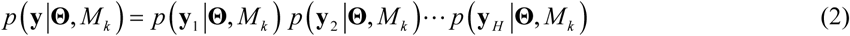

Substituting equation (2) into (1), we obtain:

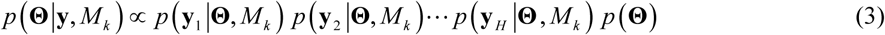

Multiplying and dividing *M-1* times by the prior probability *p*(**Θ**) in equation (3) leads to:

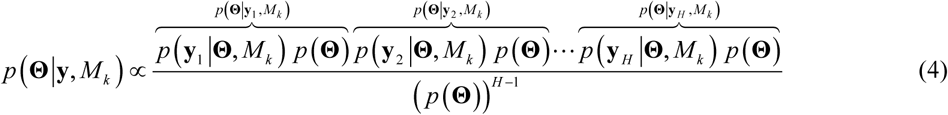

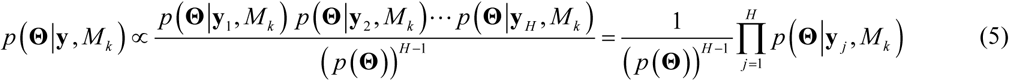

Then the posterior probability of the parameters **Θ** for a specific model *M_k_* at the federation node is expressed as a product of the posterior probabilities coming from the ‘*H*’ Hospitals. The maximum of the posterior distribution for **Θ** is achieved at 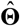 values.

#### 2.1 General Bayesian computational framework: Streaming approach

In many situations, providing a federated estimate of the parameters of interest when all local estimates are ready to use is very inefficient. Such situations become more critical when there are long delays between local parameter estimations. Hence it has been proposed, when it is tractable, the use of a subset of local estimates to obtain intermediate federated results.

Therefore as a new local estimate arrives to federation node an update of the federated estimation is performed, without recalculating parameters over again. This general scheme (Figure 2) is known as ‘Streaming’ (Broderick et al., 2013), and under the Bayesian formalism can be expressed through the following equations:

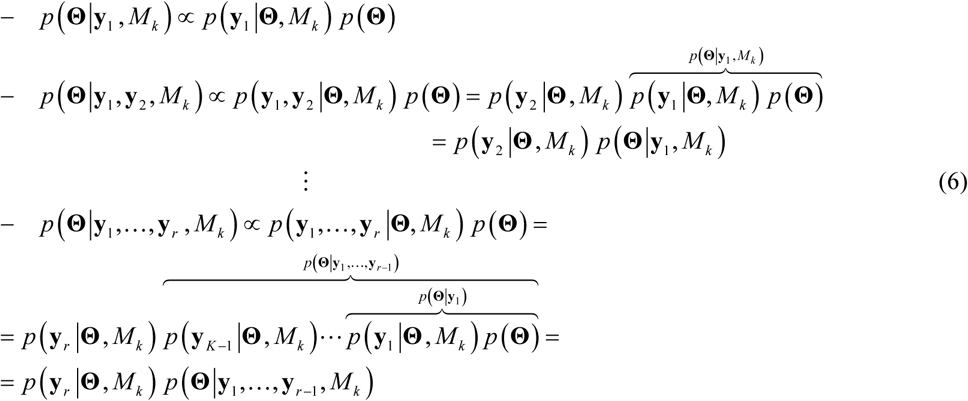

**Figure 2.**
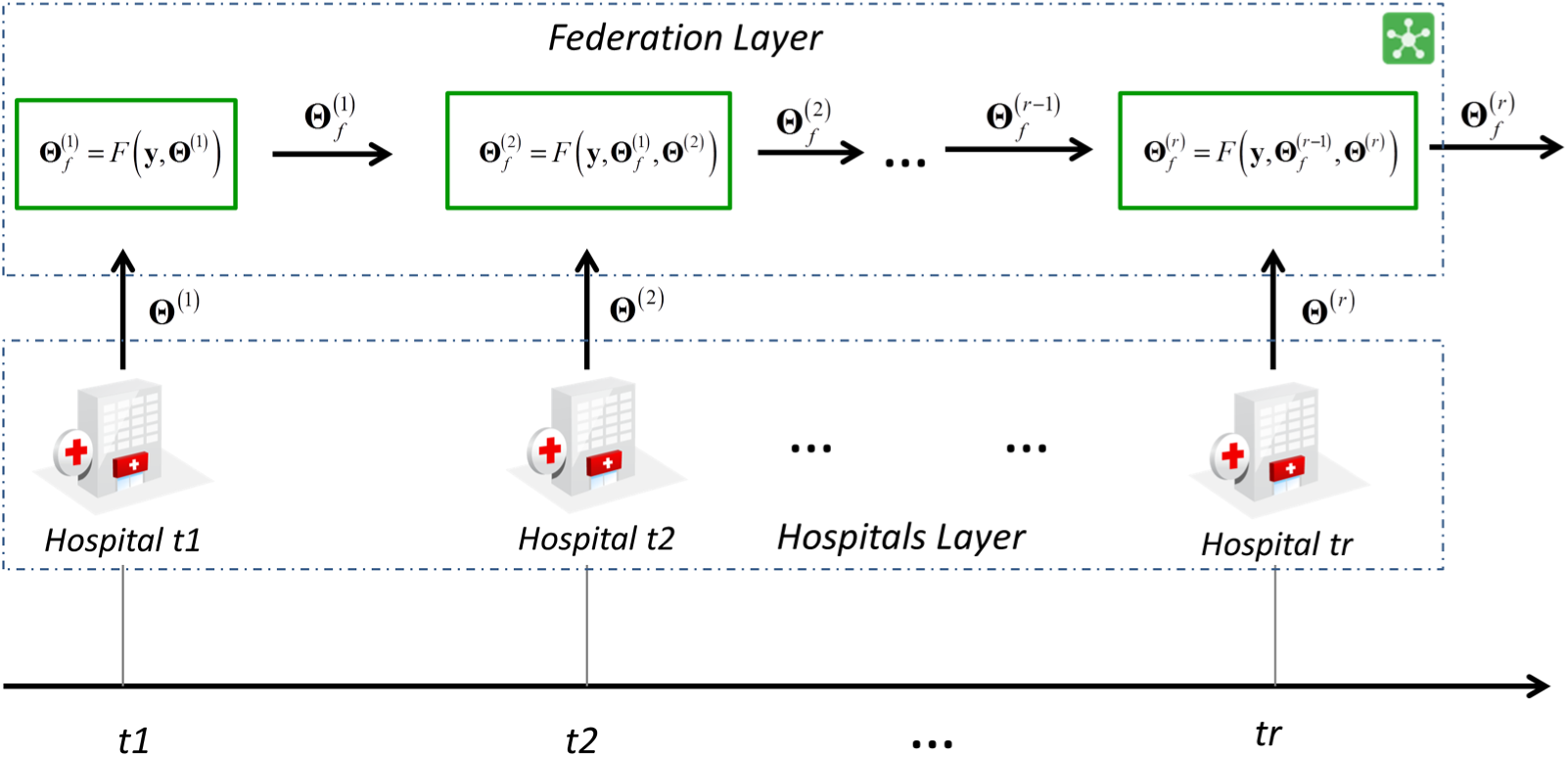
General scheme of the streaming paradigm in the Medical Informatics Platform (Human Brain Project). At the federation layer the new local estimates coming from the hospitals layer are combined with previous ones to deliver federated results.

From the recurrence equations in (6) we have that a new posterior at the federation node, when a specific ‘*r*’ data is ready from a local site, will be the multiplication of the previous posterior distribution by the likelihood function of the new data. Since the data cannot be distributed out of local sites, the posterior 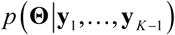 should be sent to the local site ‘*r*’ for updating the federated posterior that is finally, in a new version, pull back to the federation node.

On the other hand, equation (6) can also be rewritten as:

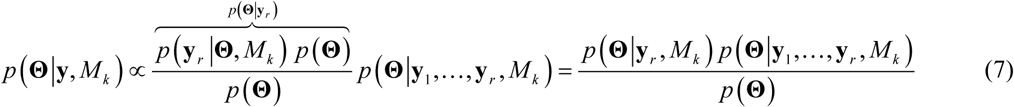

Therefore the new posterior can in addition be computed by equation (7) passing the posterior of parameters **Θ** from the ‘*r*’th hospital to the federation node to be accordingly combined with the previous posterior. That way is more convenient since the communication between federation and local nodes is preserved in only one direction, which turns out in practice to be simpler to implement.

#### 2.2 Model selection and model averaging: general definitions

As we stated at the beginning of the THEORY section, the selection of one or a set of models that explain our data is key in data analysis. In this subsection we treated the equations for the third level of inference in the Bayesian formalism known as model selection and model averaging (Hoeting et al., 1999; MacKay, 1992; Penny et al., 2006) applied to our particular case.

Given a set of ‘*K’* candidate models *M*_1_,…, *M_K_* for explaining our data, we need to calculate the posterior probability of the models to ultimately either select the best one or apply a model averaging step. In our case we assume the model selection/model averaging to be performed at the federation node, though, this can be implemented at each local site to finally be combined at the federation node. If we assume the data is generated by the same model *M_k_* in all local sites we are in the presence of a Fixed Effects (FFX) analysis. Otherwise if a more relaxed and general assumption is adopted, allowing for the possibility that different data sites use different models, a Random Effect (RFX) analysis is carried out. In the following subsections we separately analyzed both types of analysis.

##### 2.2.1 Model selection: Fixed Effects (FFX) Analysis

The probability distribution for specific model ‘*r*’ is expressed by the well-known equation:

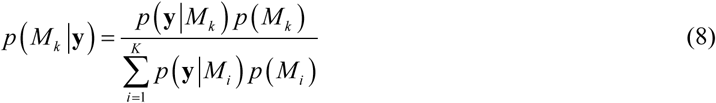

where *p*(**y**|*M_k_*) is the evidence of the model or the marginal likelihood that accounts for all data chunks associated with local sites, *p*(*M_k_*) the prior probability of *M_k_* that expresses our belief or prior knowledge about the occurrence of the model *M_k_*. The evidence of the model *M_k_* is expressed by:

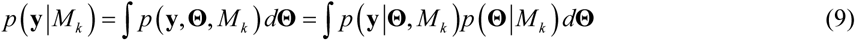

Because of the condition of independence across data sites expressed in equation (2) we have that *p*(*M_k_*|**y**) is:

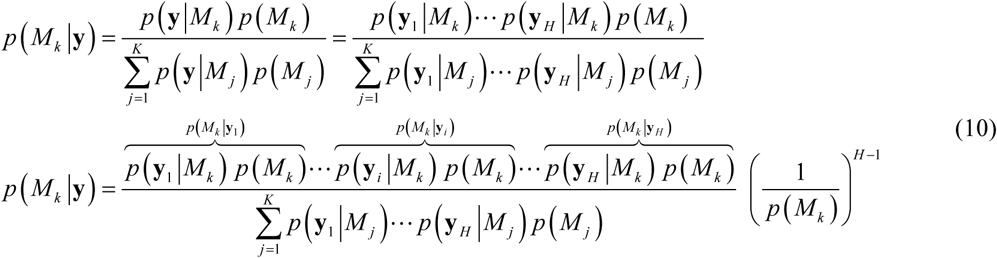

From equation (10) the posterior probability distribution of a specific model *M_k_*, at the federation node is proportional to the product of the probabilities of this model from all local data sites.

In order to select the model at federation node, that best suits the data across all local sites, we made use of the Group Bayes Factor function as has been proposed by other authors (Stephan et al., 2007):

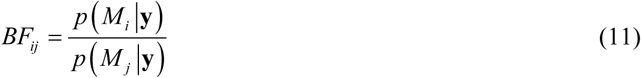

Assuming uniform model priors *p*(*M*_1_) = *p*(*M*_2_) = … = *p*(*M_K_*) = *cte* equation (11) turns:

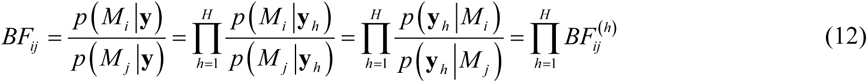

where 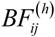 is the Bayes factor for the local data site ‘*h*’:

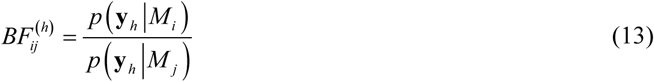

Group Bayes factor encodes the relative probability that the data were generated by one model relative to another assuming that all local data were produced by the same model. It’s assumed in practice that for *BF_ij_* > 20 provides strong evidence in favor of model *M_i_* over *M_j_*.

##### 2.2.2 Model selection: Random Effects (RFX) Analysis

In this subsection we assume the local sites use different models to fit the data that defines a Random Effect (RFX) analysis. RFX accounts for the presence of different underlying mechanisms that explain the data. This approach has been proven (Penny et al., 2010; Stephan et al., 2009) to outperform the classical FFX since its markedly more robustness in the presence of outlier data.

We adopted the methodology developed in Stephan et al. (Stephan et al., 2009) and later extended for model averaging (for family of models) by Penny et. al. (Penny et al., 2010). The proposed RFX approach determines the probability density from which the models that generate the data are sampled at different local sites.

Given the ‘*K*’ candidate models, we define a labeling process binary variable or model switch that helps us to calculate the probability that a specific hospital data was generated by the model *M_k_*. Hence, matrix S is defined as:

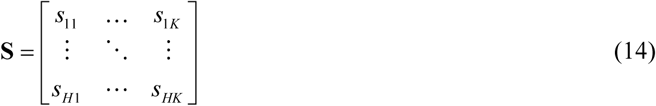

where ‘*K*’ is the number of models and ‘*H*’ number of data sites.

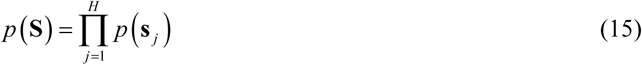

The row vector 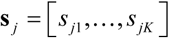 expresses the indicator variable to pick one of the ‘*K*’ models for the hospital ‘*j’*. We assume a priori that the distribution of this variable is independent across local sites, and *p*(**s***_j_*) is in general a Multinomial distribution so that:

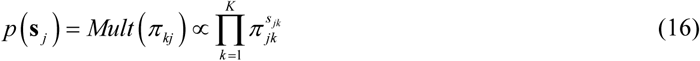

In our case 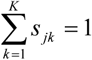 that guarantees that only one model will be selected reducing the prior to a Categorical distribution. The element *π_jk_* is the prior probability the data in hospital ‘*j*’ was generated by the model ‘*k*’. Therefore the prior probability of **S** is finally expressed as:

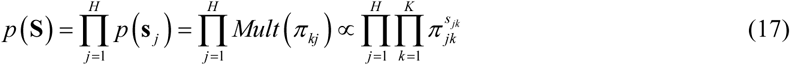

The model frequency variable represented by ***π*** = [*π*_1_,…, *π_K_*] expresses the frequency of each of the ‘*K*’ models, where 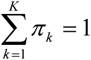. We assume that *p*(***π***) is a Dirichlet distribution so that:

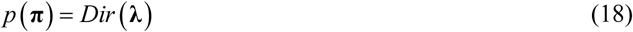

with **λ** = [*λ*_1_,…, *λ_K_*]. If a symmetric Dirichlet distribution is assumed, then *λ*_1_ = *λ*_2_ = *λ*_3_… = *λ_K_* = *λ*_0_ and:

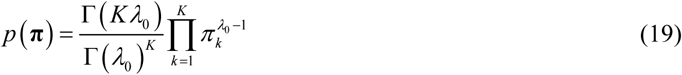

The posterior probability *p*(**π**|**y**,*λ*) provides the information required to perform model selection or model averaging at the federation node, accounting for model variability across local sites.

We used the Variational Bayes approach (see later in next subsections) to approximate *p*(**π**|**y**,*λ*), which is based on the model evidence shown in equation (9). Further details about the derivation of these equations, and an algorithm implementation for estimating the parameters of interest, can be found in (Stephan et al., 2009). Hereafter, only the final equations will be provided.

*For model indicator variable* **s**:

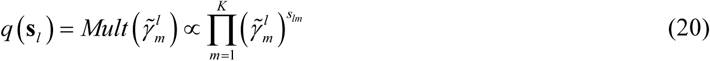

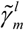 is the mean of the multinomial distribution. This is the expected number of times the model ‘*m*’ is responsible of generating data ‘*l*’. The expression for 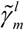 is the following:

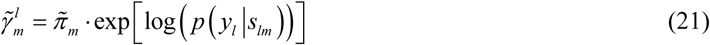

where 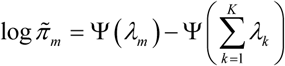, ψ(·) is the digamma function. Finally, 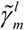 is normalized to obtain 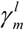 that is used in the rest of parameter estimators:

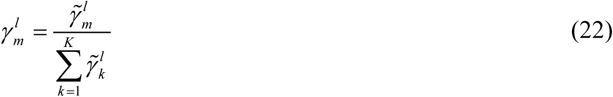

*For model frequency* **π**:

The posterior *p*(**π**|**y**,*λ*) is approximated by *q*(**π**|**y**), which takes the form of a Dirichlet distribution:

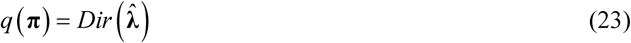

where 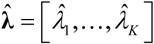, with 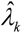 defined as:

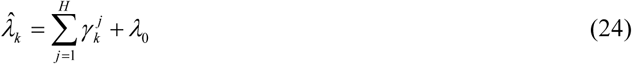

Then 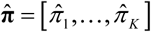, with:

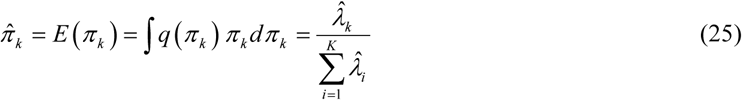

The model that best explains the overall data is defined as that with highest probability 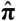, according to equation (25). The probability of model ‘*k*’, in a RFX analysis at the federation node, is expressed by:

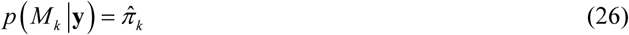

The most probable model is obtained by:

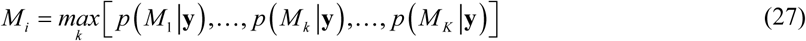

Having estimates of the model probabilities accounting for all model evidences and occurrences (at the federation node) allows us to compare two models, rank them or compare families of models. The last use-case is especially important when comparing model families of different natures.

##### 2.2.3 Model averaging

For models of a similar nature, the marginal distribution of the parameters **Θ** that accounts for the uncertainty of selecting a specific model at the federation node is given by:

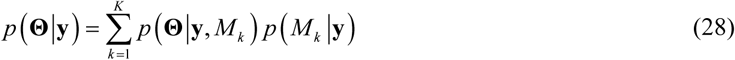

The posterior probability of model *k* at the federation node *p*(*M_k_*|**y**) may be calculated using a FFX - equation (10) - or RFX - equation (26) - analysis. Posterior probabilities of the parameters *p*(**Θ**|**y**,*M_k_*) for a specific model would be provided either by the equation (5) or (7).

### 3. Multiple Linear regression (MLR) for distributed data: probabilistic generative model

In this subsection we apply the previous general approach to the special case of the multiple linear regression model (MLR). It allows comparing different MLRs as considering subsets of explanatory variables that best explain the dependent data. This approach accounts for distributed data where only aggregates can be passed to the federation node to ultimately estimate the distribution of the parameters that best explained the data across local data sites.

Multiple linear regression allows to study the relationship between a dependent variable **y** and the explanatory variables (or independent variables) denoted by **A**. The general problem is to solve the standard linear equation posed as following:

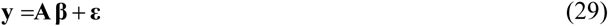

**y**: vector of dependent variable *p*×1; **A** = [*A*_1_,…, *A_N_*]: matrix of regressors or design matrix so that 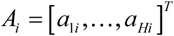, *p*×*N*; **β**: vector of regression coefficients *N*×1; **ε**: vector of experimental noise/error, *p*×1.

We have the special situation in the MIP that data **y** and **A** are naturally split and distributed over *H* different sites (hospitals). Additionally the data cannot be fetched to a federation site to solve the equation (1). Thus we have to adapt classical methods for estimating **β**, which requires combining aggregates from hospitals data.

We propose to use the Variational Bayes approach (VB) in order to solve MLR equation (29) at each local site. VB maximizes the negative ‘Free energy’ (NFE) as a surrogate measure of the log evidence or marginal log likelihood. Using either parallel or streaming paradigms, the MLR model parameters distributions and model evidences are properly combined in the federation node to provide global model parameters distributions, which account for the heterogeneity of data across local data sites.

#### 3.1 MLR equations for local sites (Hospitals): Hierarchical Bayesian Model

The MLR equation (29) is solved at each local data site. Therefore we omitted the sub index referring to a specific data site. The equations derived in this subsection are applied in the same way for each local site. In the following a VB formalism will be developed for MLR, partially treated by previous authors (Tzikas et al., 2008). The variables’ dependences of our MLR formalism is represented via the Directed Acyclic graph (DAG) shown in Figure 3.

**Figure 3.**
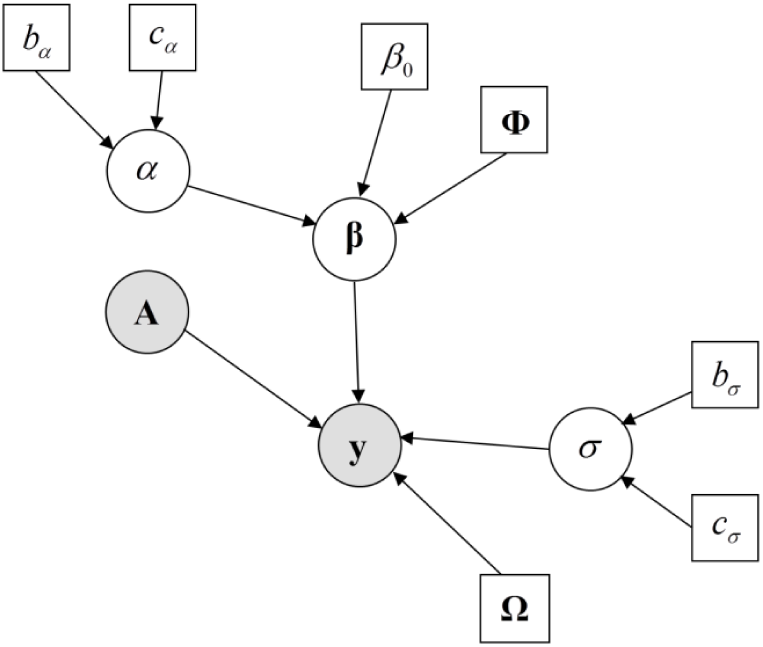
Directed Acyclic graph (DAG) of the MLR. The variables shown in squares are defined a priori. Shaded circles represent observed data whereas not shaded circles represent the model parameters (random variables) to be estimated.

The joint distribution of the data and model parameters for a specific model *M_k_* is given by:

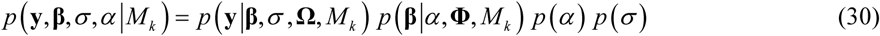

Where the vector of model parameters is **Θ** = [**β**, *σ*, *α*]; so that *σ* and *α* are the data and linear coefficients **β** precisions respectively.

A specific model *M_k_* is defined as a subset of explanatory variables. Among models the full set of variables is included whereas the absence of any explanatory variable (null model) is excluded. Thus the maximum number of models is *K* = 2*^N^* − 1. Hence, matrix A changes the number of columns for each model. Table 1 shows an example of *N*=3 explanatory variables and the seven possible models.

**Table 1.** Example of the MLR definition for three explanatory variables case. The ones in the table indicate that specific variables are included in the model. In this case we have *N*=3 variables with *K*=7 possible models.

| | $\beta_1$ | $\beta_2$ | $\beta_3$ | MLR Equation |
| --- | --- | --- | --- | --- |
| $M_1$ | 0 | 0 | 1 | $y = A_3 \beta_3$ |
| $M_2$ | 0 | 1 | 0 | $y = A_2 \beta_2$ |
| $M_3$ | 0 | 1 | 1 | $y = [A_2 \ A_3] \begin{bmatrix} \beta_2 \\ \beta_3 \end{bmatrix}$ |
| $M_4$ | 1 | 0 | 0 | $y = A_1 \beta_1$ |
| $M_5$ | 1 | 0 | 1 | $y = [A_1 \ A_3] \begin{bmatrix} \beta_1 \\ \beta_3 \end{bmatrix}$ |
| $M_6$ | 1 | 1 | 0 | $y = [A_1 \ A_2] \begin{bmatrix} \beta_1 \\ \beta_2 \end{bmatrix}$ |
| $M_7$ | 1 | 1 | 1 | $y = [A_1 \ A_2 \ A_3] \begin{bmatrix} \beta_1 \\ \beta_2 \\ \beta_3 \end{bmatrix}$ |

#### 3.2 Likelihood

We assumed **y** is normally distributed as follows:

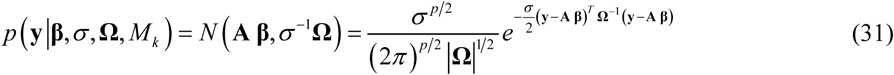

The experimental error (noise) is considered to have a normal distribution so that:

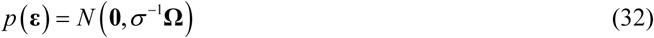

where *σ* is the error precision; **Ω** is the covariance matrix modeling the correlation structure present in our data.

#### 3.3 Prior distributions

This subsection provides the mathematical form of the prior probability distributions assumed for the parameters of interest.

*Regression coefficients* **β**:

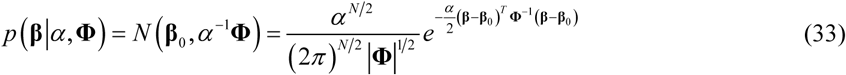

*Precision of the regression coefficients α:*

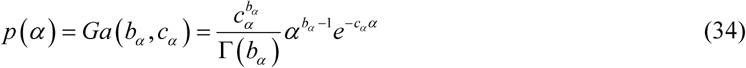

*Precision of the likelihood function σ:*

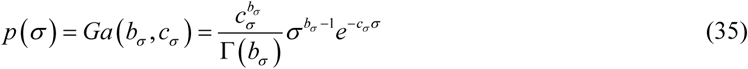

#### 3.4 Posteriors

Our aim is to estimate the posterior distributions of the unknown parameters **Θ** = [**β**, *σ*, *α*] and the evidence of the different models *M*_1_,…,*M_K_*. In doing so, we use the *Variational Bayes* formalism (VB) (Beal, 2003; Lappalainen and Miskin, 2000). This is an efficient method already used in previous publications (Penny et al., 2005; Trujillo-Barreto et al., 2008). It is based on the Variational Free Energy method of Feynman and Bogoliubov, which provides an approximate factorized posterior distribution *q*(**Θ**|**Y**), with minimal Kullback–Leibler (KL) divergence from the true posterior *q*(**Θ**|**y**) = *p*(**β**,*σ*,*α*|**y**).

We consider the following factorization of the approximate posterior distribution:

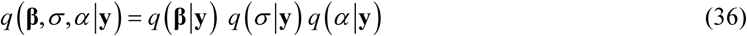

The following subsections provide details about the derivation of the approximate posteriors for each parameter. In general, the approximate posterior *q*(**Θ**|**Y**) for the *i*-th subset of parameters is expressed by *q*(*θ_i_*|**Y**) as:

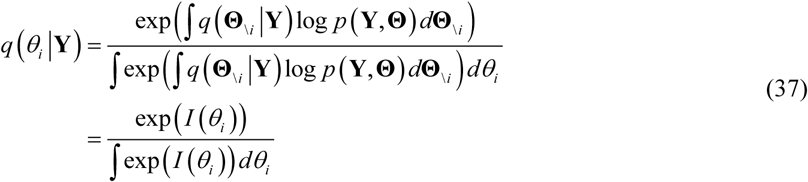

where 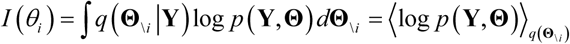, and **Θ**_\i_ indicates parameters not present in the *i*-th group of parameters.

##### 3.4.1 *Posterior for regression coefficients* β

The approximated posterior distribution of *q*(**β**|**y**) the regression coefficients **β** is expressed by:

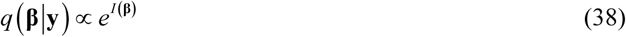

where

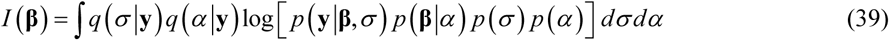

Finally *q*(**β**|**y**) is a normal distribution so that,

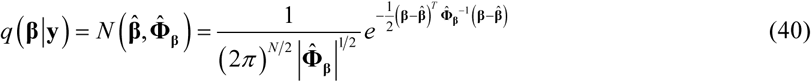

The mean and covariance matrix of this multivariate normal distribution is given by:

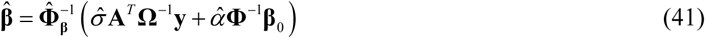

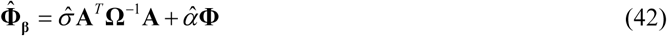

##### 3.4.2 Posterior for the precision of the regression coefficients α

In this case the approximated posterior distribution for α is obtained by:

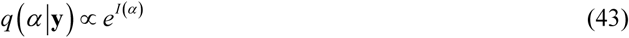

where

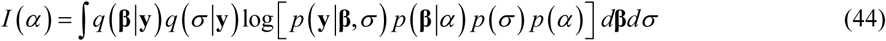

After some algebra solving equation (44) we see that the posterior distribution of *α* is approximated by a gamma distribution so that:

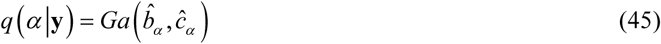

where

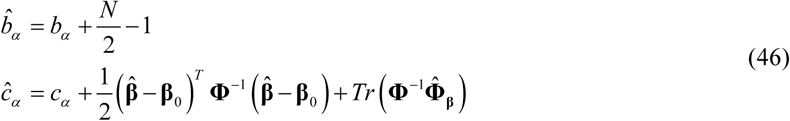

Finally, the expected value of *α* is:

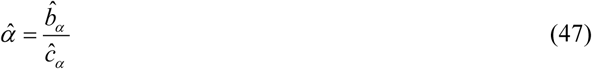

##### 3.4.3 Posterior for the precision σ

The approximate posterior distribution for *σ* is obtained by:

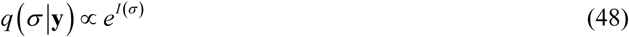

where

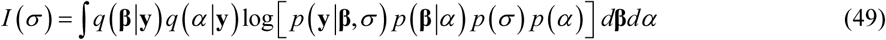

After some algebra, we obtain the following gamma distribution:

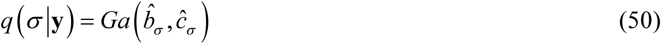

where

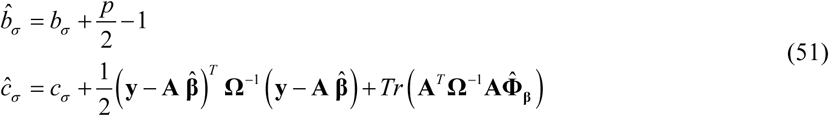

The expected value of *σ* is given by:

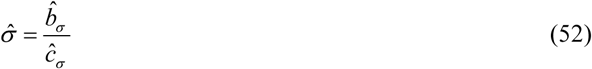

#### 3.5 Free Energy and KL divergences

The free energy function is expressed as:

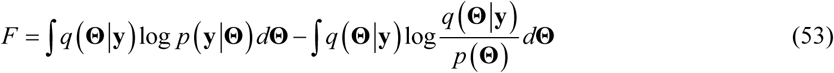

where the first term 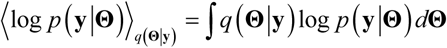 is the mean likelihood over *q*(**Θ**|**y**) and 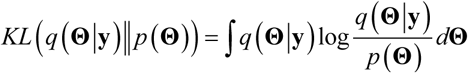 the Kullback Leibler divergence between the approximated posterior and prior distributions.

After some algebraic work, the expectation of the Likelihood is given by:

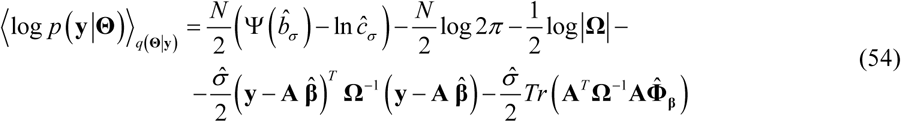

where *Tr*(·) is the trace operator.

Similarly, the Kullback Leibler divergence between the approximated posterior and prior distributions is expressed as:

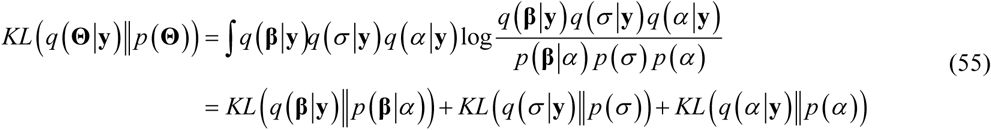

After some algebraic operations we obtain:

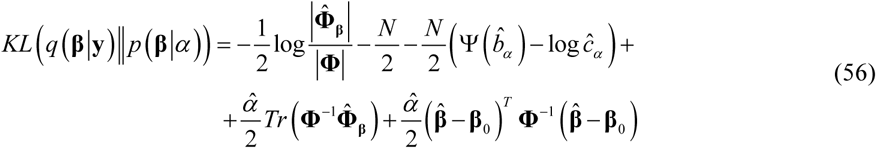

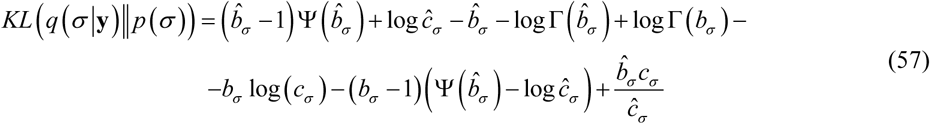

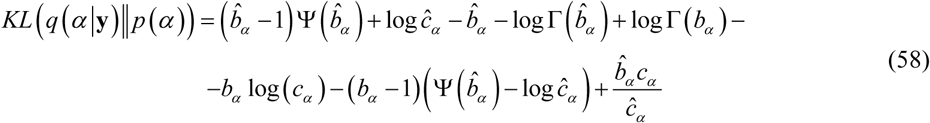

Equations (54), (56), (57) and (58) provides the necessary quantities to evaluate the Free energy function. The dependence among the posteriors of the model parameters **Θ** = [**β**,*σ*,*α*] in equations (41), (46) and (51) leads to an iterative algorithm. The Free Energy is evaluated at each iteration, and is expected to continue to increase until the convergence criterion is met. This is defined as 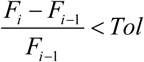, where *F_i_* and *F_i_*_−1_ are the free energy at iteration ‘*i*’ and ‘*i*-1’ respectively, *Tol* is the tolerance with values defined in 0 ≤ *Tol* ≤ 1. Since the Free Energy is a surrogate measure of the log model evidence, the model probabilities - equation (8) - at local nodes, are given by the expression:

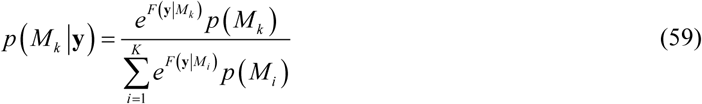

where the model evidence is expressed by:

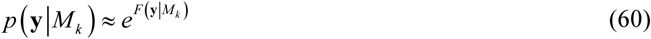

As a final step, the model evidences and the distributions of the parameters **Θ** = [**β**,*σ*,*α*] from each local node are submitted to the federation node to be properly federated.

The general iterative algorithm that runs at each local node is the following:

###### Local Node –General Algorithm

- Local data **y** and **A** assessment.
- Definition of the stopping criteria tolerance ‘*Tol*’
- Definition of **Ω**, **Φ** as the covariance structure for **y** and **β** respectively.
- Initialize hyperparameters: *b_σ_*, *c_σ_*, *b_α_*, *c_α_*
- For *k*=1 to *K* (Number of Models)

- Iteration ‘*i’*

- Compute 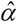 using equation (47)
- Compute 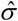 using equation (52)
- Compute 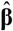 and 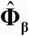 using equations (41) and (42) respectively
- Compute Free Energy *F_i_* for iteration ‘*i’* combining equations (54), (56), (57) and (58) Repeat until 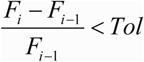
- Evaluate evidence of model *M_k_* using equation (60)
- End (Number of Models)
- Send to the Federation node: evidences of the ‘*K*’ models and distributions of the parameters 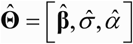

#### 3.6 MLR equations for the federation node

At the federation node, the *MLR* is estimated accounting for parameter distributions and model evidences coming from local nodes. In order to obtain expressions (see next subsection) for the federated *MLR* model parameters we made use of the general equations developed in all subsections 2.x.

##### 3.6.1 *Linear model coefficients* β

At each local node, this parameter has a normal distribution given by equations (40), (41) and (42). The distribution of **β** at the federation node using equation (5) is given by:

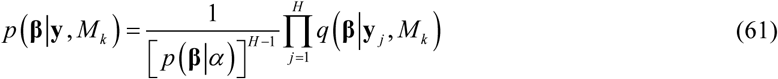

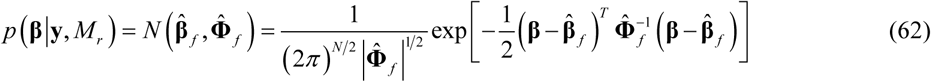

where,

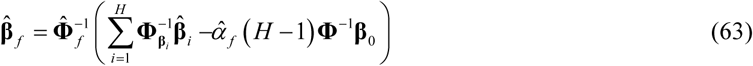

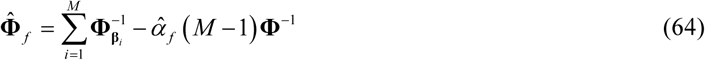

##### 3.6.2 Precision of the regression coefficients α

At each local node, *α* follows a Gamma distribution given by equations (45), (46) and (47). After some algebraic operations on equation (5) we see that *α* also distributes as gamma at the federation node so that:

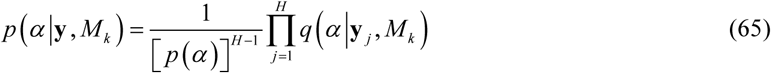

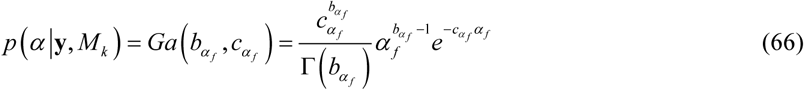

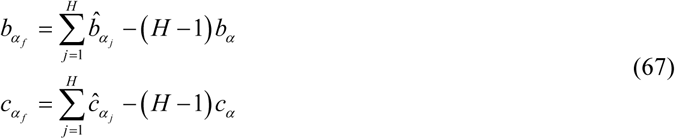

The expected value for *α* is expressed by:

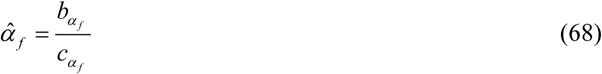

##### 3.6.3 Precision of the likelihood function σ

At each local node, *σ* has a Gamma distribution given by equations (50), (51) and (52). After combining the local estimates through equation (5) this model parameter distributes as gamma at the federation node. This probability distribution is expressed as:

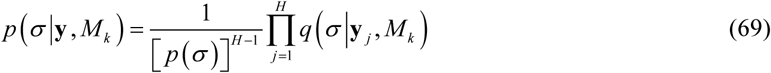

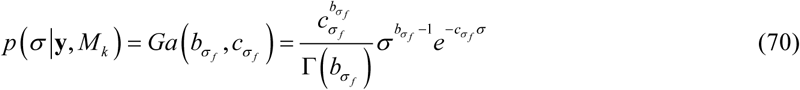

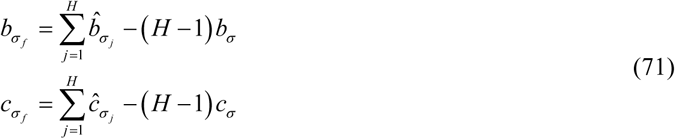

with the expected value given by:

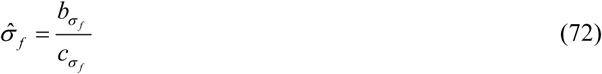

##### 3.6.4 Model selection and averaging for MLR at the federation node

As we mentioned before, a specific model *M_k_* in the MLR is defined as a subset of explanatory variables (see Table 1). This means we are looking for a subset of *β_i_* in **β** that best contributes explaining the data across hospital sites. The number of possible models ‘*K*’ is *K* = 2*^N^* − 1, taking into account that each explanatory variable can be present or not. The model selection at the federation node is conducted by using either a FFX or RFX approach given by equation (10) or (26) respectively. The model averaging step, via expression (28), provides the estimation of the parameter distributions irrespective of the model selected. For the regression coefficients this equation takes the mathematical form:

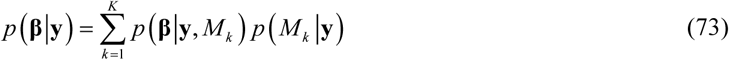

The general algorithm that runs at the federation node is the following:

###### Federation Node –General Algorithm

- Fetch from the ‘*H’* local nodes the local parameter expectations: 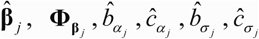 for all models *M_k_*, *k* = 1,…, *K*.
- Estimation of the federated expected value 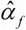 of the precision of the linear model coefficients using equation (68)
- Estimation of the federated expected value 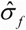 of the precision of the data distribution using equation (72).
- Estimation of the federated linear coefficients 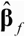 and the covariance matrix 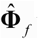 using equations (63) and (64) respectively.
- Execution of the Model selection using a ‘*Random Effects (RFX) Analysis’* (see the corresponding section in the text).
- Model averaging using equation (73).
- Submit to the user the distributions and its expectations 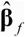 for the model parameters and the model that best explains the data (set of explanatory variables that best explains the data).

## CONCLUDING REMARKS

In this paper we presented a Bayesian formalism to deal with distributed, highly heterogeneous and big data, based on previous theoretical developments. This formalism is part of the Bayesian computational core embedded in the Medical Informatics Platform (MIP) of the European Human Brain Project (www.humanbrainproject.eu). As an application example we applied the general approach outlined in this work to the multiple linear regression (MLR), which is considered one of the workhorses of statistics in the neuroimaging field. Though the adaptation of the MLR to a distributed environment might be considered straightforward, in this work we extended it with a model selection and averaging approach. This is crucial for selecting appropriate models to explain highly heterogeneous data.

The federation node is defined as the site in which the local aggregates quantities are combined to deliver a final result accounting for the diversity of the different hospital data (local nodes). We provided approximated probability distributions of the MLR parameters using the Variational Bayes formalism (VB). VB approximates the parameters probability distributions based on the minimization of the negative free energy (NFE) as a surrogate measure of the log evidence. NFE enables the estimation of model probabilities to ultimately performing the model selection and averaging step at the federation node. In this respect, we brought into our formalism the methodology proposed by Stephan et al. (Stephan et al., 2009) (for ‘*Dynamic Causal Modeling*’-DCM-) to account for the heterogeneity of the possible mechanisms that explain data at different hospitals. This is an important aspect in order to integrate highly heterogeneous data coming from different subjects and hospitals across the globe. For the case of a large number of explanatory variables (*N* ≫ 1) an exhaustive evaluation of all model evidences is computationally prohibitive. In such cases a Markov Chain Monte Carlo (MCMC) approach ought to be adopted. MCMC would sample the mass of the models probability distribution allowing, as before, model selection and model averaging steps to be conducted. In addition, a factorization over **β** coefficients in the VB scheme -equation (36)- should be assumed at local nodes in order to get tractable estimations.

The implementation of our theoretical formalism and its application, in both simulated and real data, will be addressed in a separate work. Its performance under different conditions and the limits of its application will be evaluated through simulated data. Several situations should be considered: a) unbalanced data across local nodes; b) influence of the number of local nodes on model selection and parameters estimation; c) robustness to the presence of outliers; d) robustness of the estimations when the number of explanatory variables increases, etc.

The present work is the first of a series of papers adapting, through Bayesian formalism, the existing statistical models to distributed and big data in order to ultimately discover underlying mechanisms of brain anatomy and function (using the Medical Informatics Platform). This approach is not limited to the Neuroscience field; it can be applied in other branches of science and technology.

## ACKNOWLEDGEMENTS

The research leading to these results has received funding from the European Union Seventh Framework Programme (FP7/2007–2013) under grant agreement No 604102 and the European Union’s Horizon 2020 research and innovation programme under grant agreement No 720270 (HBP SGA1). BD is supported by the Swiss National Science Foundation (NCCR Synapsy, project grant Nr 32003B_159780 and SPUM 33CM30_140332/1), Foundation Parkinson Switzerland and Foundation Synapsis.

